# NaviGraph: A graph-based framework for multimodal analysis of spatial decision-making

**DOI:** 10.1101/2025.05.18.654725

**Authors:** Amit Koren Iton, Elior Iton, Lior Bikovski, Daniel M. Michaelson, Pablo Blinder

## Abstract

Understanding spatial decision-making requires interpreting multimodal data streams through a unified analytical lens that relates them to one another and to the layout of the underlying task. We developed NaviGraph (Navigation on the Graph), an open-source framework that formalizes spatial tasks as graphs to provide a topological scaffold for data integration. This approach solves the challenge of aligning disparate data streams while standardizing spatial analysis across diverse laboratories and behavioral paradigms. NaviGraph transforms decision points into nodes and paths into edges, enabling the computation of both conventional and graph-based metrics, such as unique node exploration and trajectory efficiency. We demonstrate its utility and enhanced sensitivity through a complex memory task, where graph-derived metrics revealed subtle recall impairments in female Apolipoprotein E ε4 (ApoE4) mice - the primary genetic risk factor for Alzheimer’s disease - that were undetectable by conventional measures alone. We further illustrate multimodal alignment by mapping retrosplenial cortex calcium imaging and head orientation onto the graph structure, offering a topological perspective on decision-point dynamics. With its modular, plugin-based architecture, NaviGraph provides a standardized environment for the exploration of multimodal data across diverse spatial paradigms.

## Introduction

Behavioral experiments are a fundamental component of preclinical animal research, crucial for understanding cognitive processes and assessing therapeutic interventions ^1,2^. These tests span a wide range of modalities, from highly controlled, head-restricted setups relying on repeated trials ^3,4^, to ecologically relevant tasks replicating real-world environments in which animals engage in unrestrained naturalistic behaviors ^5–7^. However, balancing behavioral complexity and natural behaviors, while ensuring feasibility in controlled laboratory settings, remains a significant challenge. For rodents, complex mazes help address this by mimicking real-world navigation through sequential decisions, supporting robust learning and memory retention ^8^. While these designs enable more naturalistic observations in laboratory settings, the most common approach to behavior analysis predominantly relies on what can be described as classic metrics, such as latency, velocity and error rates ^9–11^. Though widely applied, these measures don’t address relational and structural properties of navigation that are critical for capturing the full complexity of decision-making, and are easily influenced by confounding factors such as stress, motivation, or physical state ^12,13^. Topological analysis, which focuses on relational rather than purely spatial properties, has shown value across diverse domains, including transportation and protein folding ^14^, but remains underexplored in behavioral research, despite its potential to complement the limitations of classic metrics. As a result, the architecture of decision-making often goes unmodeled, and behavior is frequently reduced to isolated events rather than interpreted as a structured process.

These conceptual limitations are further accentuated by the increasing complexity of modern behavioral setups, which now enable simultaneous collection of multiple parameters during experiments, including neurophysiological ^15,16^, behavioral ^17,18^, and physiological ^19^ data streams. While these innovations have greatly enhanced our ability to study cognition, the diverse nature of these datasets presents significant obstacles ^20,21^ for the analysis and later sharing of study results. Though tools that synchronize behavioral and neuronal data have started to emerge ^22^, they mostly focus on enriching neuronal analysis perspectives in simple behavioral setups ^23^, whereas complex spatial designs enabling trajectory-based metrics receive limited attention. As such, there is currently no standardized open-source tool capable of seamlessly integrating diverse data streams into a unified analysis framework while being adaptable enough to encompass a wide range of spatial decision-making behavioral setups.

To address these gaps we developed *NaviGraph* (Navigation on the Graph), a flexible pipeline designed to function as a universal integration point of diverse data streams within a graph-based framework. At the core of NaviGraph is a topological representation of the behavioral arena, where decision points are modeled as nodes and the paths between them as edges. Importantly, these decision points can be implicitly defined by the layout of the behavioral apparatus (such as T, Y or more complex mazes), arbitrarily defined by the experimenter or inferred from observed behaviors. With the synergistic combination of classic and topological approaches, NaviGraph provides a structured yet flexible analysis framework, enabling the integration of multiple data streams and parameters into a cohesive multilayer graph ^24^. By allowing data to exist within their own layer, while maintaining connectivity between layers through the same graph structure, interactions between diverse data at each decision point can be captured in a straightforward manner, enabling experiment-level interpretation. The pipeline includes a graphical user interface for easy setup, as well as various modules for data processing, computation of classic behavioral and graph-based metrics, multi-stream data alignment, aggregation, and visualization of results. We adapted a plug-in-based software structure providing researchers with complete freedom to extend NaviGraph with custom data types or workflows. As we demonstrate, this approach enhances the detection of subtle behavioral nuances, many of which may remain obscured when using traditional or single-stream analysis.

We applied NaviGraph within an automated, trial-based version of the complex binary maze introduced by Rosenberg et al. ^8^, and implemented a robust learning and memory task, emphasizing minimal human intervention. By leveraging both classic and topological measures, we show how these complementary approaches provide a deeper understanding of mouse behavior in these types of environments, both under normal and pathological conditions. We further demonstrate the platform’s multi-stream capabilities by integrating head orientation and neuronal activity from unrestrained calcium imaging recordings with miniaturized microscopes ^25^, in order to showcase potential for future applications. Overall, NaviGraph’s approach provides a valuable tool that leverages the power of graph theory to offer a standardized, flexible and sensitive analysis of spatial navigation and decision-making, enabling multimodal behavioral neuroscience without requiring advanced computational skills or custom infrastructure.

## Results

### NaviGraph pipeline structure

NaviGraph solves a critical challenge in behavioral neuroscience, namely standardizing and integrating outputs from multiple sources across multi-session experiments into a single data-structure suitable for decision-making analysis across these modalities. Through its plugin-based architecture, the system functions as a universal integration point, capable of linking pose tracking, neuronal recordings, physiological parameters, and user-defined measures within a single analytical framework. NaviGraph’s workflow is illustrated in Figure 1, which describes the structure of the pipeline and its capability to unify data streams and behavioral parameters within a biologically meaningful graph-based framework.

**Figure 1:**
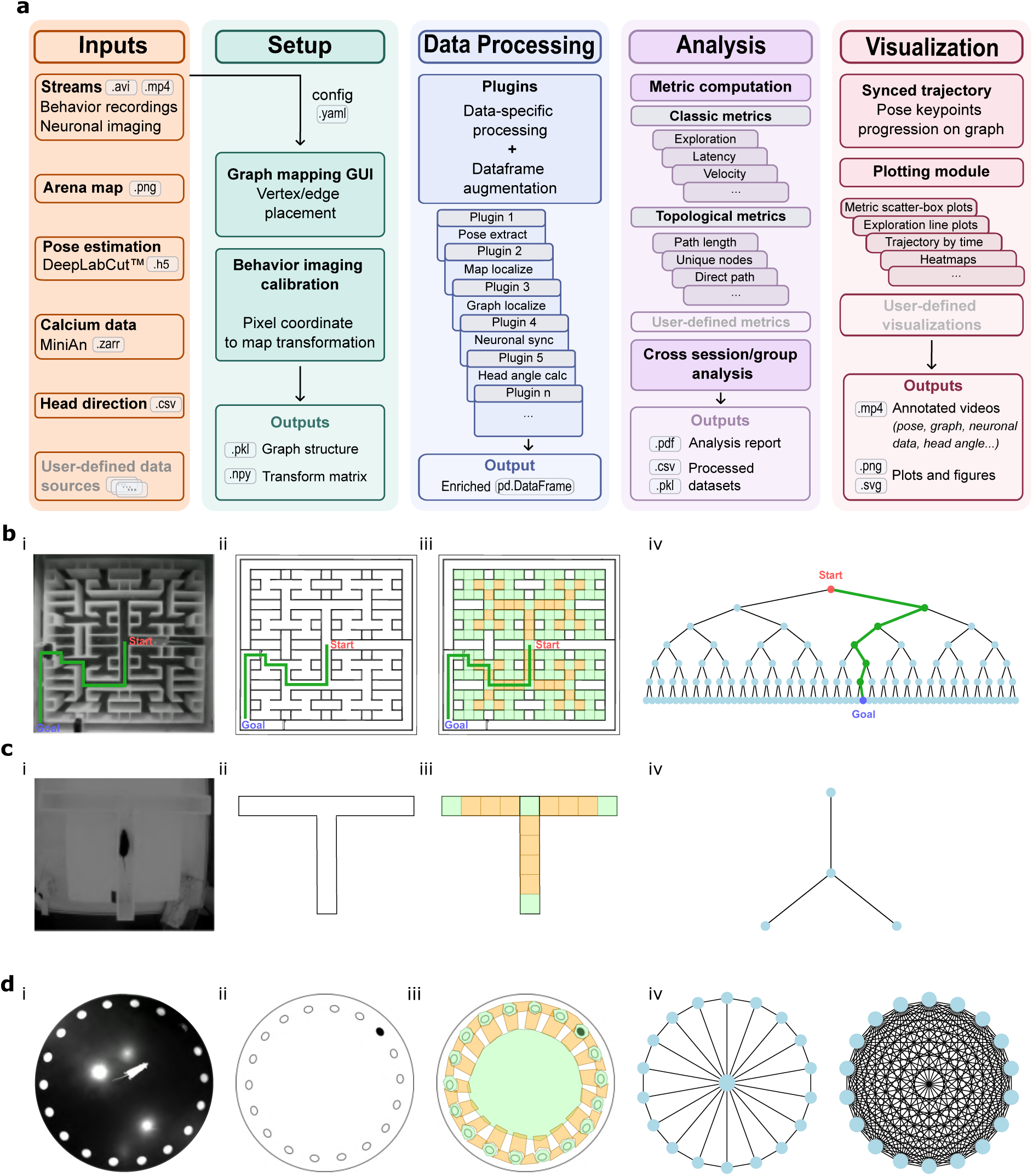
NaviGraph workflow and adaptability across spatial behavior paradigms. (a) Pipeline overview. Input streams include behavioral recordings, neuronal imaging, pose estimation, calcium activity, head orientation, and additional user-defined data. Setup comprises environment-to-graph mapping via a graphical interface and calibration of pixel to spatial coordinates. Data processing is plugin-based, producing a unified dataset for analysis. Modules compute classic and topological metrics, with support for user-defined measures and cross-session comparisons. Visualization generates synced trajectories, graph overlays, and quantitative plots. (b–d) Pipeline paradigm flexibility examples. (b) Seven-level deep binary maze: raw image (i), schematic (ii), region segmentation (iii), and corresponding graph (iv). (c) T-maze: raw image (i), schematic (ii), region segmentation (iii), and graph (iv). (d) Barnes maze: raw image (i), schematic (ii), region segmentation (iii), and two out of the many optional graph abstractions for this specific environment (iv).

Initially, pose estimation datasets are spatially registered to the maze layout through the calibration and validation plugin, that ensures accurate transformation of raw pixel coordinates into topological representations, namely, nodes and edges of the graph structure. This enables precise mapping of the trajectory onto the graph, which serves as a common scaffold for all subsequent analysis. Within this framework, NaviGraph has built-in plugins that compute both conventional behavioral metrics (e.g., time from a to b and velocity) and graph-based topological metrics (e.g., number of unique nodes traveled and number of nodes on direct path to reward). If available, additional data streams such as neuronal activity and head orientation are synchronized and coupled to the same graph, enabling node and path specific multimodal analysis.

A defining feature of NaviGraph is its ability to traverse data across domains. The same experiment can be interrogated in the temporal domain (time-series dynamics), the spatial domain (coordinate-based trajectories), and the graph domain (topological structure), providing complementary perspectives on behavioral and neuronal function. Moreover, outputs enable aggregation across trials, sessions, and groups, supporting longitudinal and cross-condition comparisons. In addition, NaviGraph has a built-in capability to run random-walker agents within the graph structure, allowing the generation of null hypotheses for behavioral performance; a feature that becomes more relevant once the complexity of the behavioral layout increases.

While many currently available tools in the field focus on isolated data modalities or require substantial post-processing, NaviGraph offers an experiment-level analysis approach. Its flexible open-source design supports a wide range of spatial layouts and experimental paradigms (Fig.1), making it a broadly adaptable, extensible, and reproducible platform for multimodal behavioral neuroscience research. A more detailed overview of the computational modules and functions is provided in the Methods section and accompanying code repository.

### Behavioral application of NaviGraph

We showcase NaviGraph’s ability to analyze behavior with a combination of classic and topological measures (Fig. 2). Our trial-based automated version of the seven-level binary tree maze paradigm ^8^, was employed to assess learning and memory in male and female C57BL/6 wild-type (WT) mice. In each trial of this task, mice were required to navigate through a series of six correct decisions from the maze start to reach a liquid reward port (Fig. 2a). Following one week of self-imposed water restriction (home-cage water made slightly sour with citric acid ^26,27^), each mouse completed a learning session which included four consecutive rewarded trials absent of experimenter intervention, followed by a single memory trial 72 hours later. To minimize reliance on spatial cues, the task was conducted in the dark, and maze rotation was applied between sessions. Potential odorant cues from the reward port were masked by placing the liquid rewarded solution around the corners of maze exterior.

**Figure 2:**
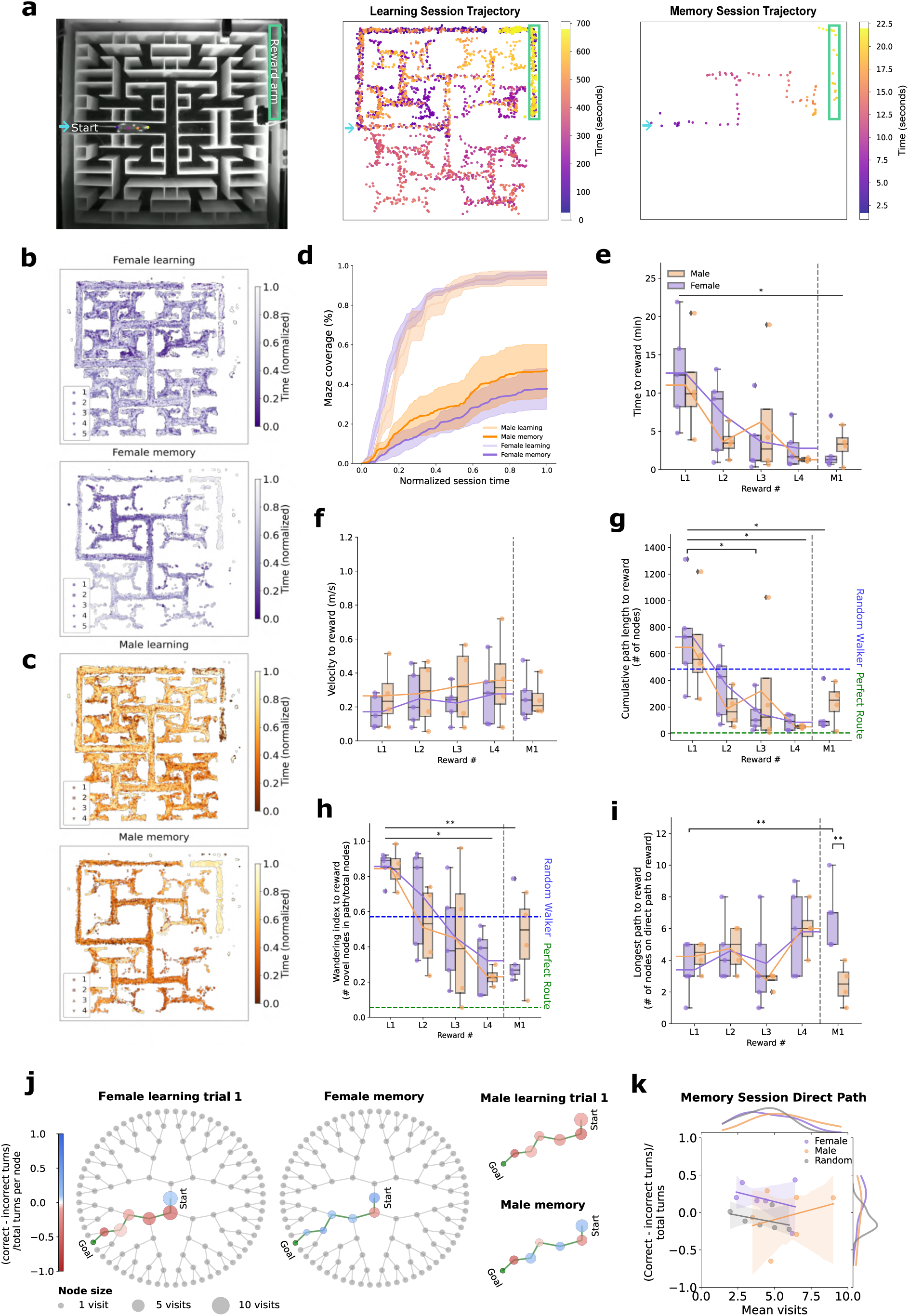
NaviGraph synergistic topological and classic behavioral analysis in male and female wild-type mice. (a) Complex maze paradigm. Left: Maze structure. Middle: Representative learning session trajectory. Right: Representative memory session trajectory. Entrance and reward arm indicated by blue arrow and green box respectively, time progression depicted by colorbar. (b, c) Trajectories over session time. Normalized trajectories of individual C57BL/6 wild-type (WT) n=5 female (b) and n=4 male (c) mice in learning (top) and memory (bottom) sessions respectively. (d-i) NaviGraph metrics. Females and males shown in purple and orange, respectively. Four consecutive learning session trials represented by rewards L1-L4; memory session represented as M1. (d) Maze coverage during normalized session time, shown as mean ± SEM. (e) Time taken to reach the reward. (f) Average velocity to the reward. (g) Cumulative path: total number of nodes passed to reach reward. Mean of 5000 random walker bouts on the graph depicted by the blue dashed line. Direct path to reward depicted by green dashed line. (h) Wandering index: proportion of unique nodes visited. (i) Longest path to reward: number of nodes passed on the direct path to the reward. Data in box plots depicted as median ± interquartile range, lines reflect group means; * p<0.05. (j, k) Confidence index and node visits shown on the binary tree representing the maze. (j) Decision confidence levels, calculated as the difference between mean correct and incorrect turns relative to the total number of turns per node, for females (left) and males (right) during first trial of learning and memory sessions. Node sizes represent mean group visit frequency to each node, and color intensity denotes the confidence index.

Time-based trajectories of both female and male WT mice during these spatial learning and memory sessions revealed distinct patterns (Fig. 2b and 2c), with an overall shift from broad exploration in the learning session to more efficient navigation in the memory. NaviGraph maze coverage metric (Fig. 2d) quantifies this feature, providing an overview of normalized maze exploration across sessions. In the memory session, we observed a trend where males explored a larger portion of the maze relative to females. Importantly, both sexes show reduced maze coverage in the memory session relative to the learning session, indicating an overall memory retention of the maze structure; even when a limited familiarization period with this new environment was provided.

NaviGraph further captures classic behavior metrics such as time taken to reach the reward and average velocity (Fig. 2e,f). The first trial of the learning session (L1) represented each animal’s initial exposure to the maze layout, noting that in order to shorten the previously reported exploratory phase ^8^, animals were introduced to the paradigm via the reward arm. We observed gradual reduction in the time taken to reach the reward across the learning trials in both sexes (Fig. 2e) (p=0.027 effect of learning trial number; two-way Repeated Measures (RM) ANOVA). In the memory session (M1), females reached the reward slightly faster than males, but no significant differences between the groups were shown. Both groups displayed significantly shorter times in the memory session relative to the first trial of the learning session (p=0.001 for females and p=0.01 for males). In both sexes there was general conservation of velocity through the trials (Fig. 2f). No significant velocity differences were observed between males and females in either session, indicating that speed does not underlie the time-based performance differences.

To contextualize mouse performance within the topological measures, we generated benchmarks for random graph navigation (i.e. a random-walk agent) by averaging 5000 random walks on the tree graph, with and without backtracking. Our results revealed that random walks without backtracking, which are allowed to turn around only upon reaching a dead end, yielded an average cumulative path length (total number of nodes visited in each trial) of 485.38 nodes ± standard deviation (SD) 490.98 nodes on its way to the reward location. In contrast, the random walks with backtracking resulted in an average cumulative path length of 1298.14 nodes ± 1365.89. When calculating the “wandering index”, referring to the number of novel nodes visited relative to the total number of nodes in the maze, there was less contrast between the backtracking (0.576 ± 0.282) and non-backtracking walks (0.567 ± 0.281). Consequently, the analysis focused on the results from the non-backtracking random walks, as these values aligned more closely with actual behaviors of the mice and provided a more realistic benchmark for evaluating performance.

We show that both cumulative path length and wandering index values of the first learning trial (L1) were, as expected, comparable to the random walk benchmark, while in the final learning trial (L4) these measures shifted towards the perfect route to the reward. Moreover, the cumulative path length (Fig. 2g) decreased significantly across the learning session, indicating improved navigational efficiency (p=0.01 for L1-L3 and L1-L4 in females, p=0.04 for L1–L4 in males; Tukey’s post hoc following 2-way RM ANOVA). Similarly, the wandering index (Fig. 2h), also declined progressively (p=0.017 in females, p=0.006 in males), further supporting the presence of spatial learning.

To assess memory retention, we compared performance in the final learning trial (L4) and the memory trial (M1) conducted 72 hours later. No significant differences were found in either cumulative path length or wandering index, indicating that mice retained spatial memory across sessions. Furthermore, comparisons between L1 and M1 demonstrated significant reductions in cumulative path length (p=0.002 in females; p=0.02 in males) and wandering index (p=0.001 in females; p=0.0006 in males). In the memory session (M1) females showed a trend toward lower exploratory behavior, suggesting a slightly more refined spatial memory recall.

To assess spatial insight of the maze structure, we analyzed the length of the direct path to the reward, defined as the sequence of consecutive correct choices leading to the reward location (Fig. 2i). A longer direct path to the reward indicates fewer mistakes, reflecting a more accurate internal representation of the maze structure ^8,28^. This measure reflects the ability to navigate the maze efficiently, suggesting a stronger cognitive map and reliance on learned spatial representations rather than exploratory or random navigation strategies ^29,30^. We observed that both sexes showed an overall positive learning curve along the trials of the learning session, which manifested as a gradual increase in the number of nodes on the direct path to the reward. Interestingly, females exhibited longer direct paths to the reward during the memory session, with fewer deviations compared to males (p=0.008; Mann-Whitney test), indicating a higher degree of maze structure retention during navigation. This is further supported by the significantly longer direct paths to the reward in females when comparing the recall session to the first trial of the learning session, in which the navigation was close to random (p=0.0021).

A key feature of NaviGraph is the ability to topologically visualize behavior on the graph, making the learning process visible on the route itself. Here, we display decision confidence and visit counts on the binary tree representing the maze (Fig. 2j,k), where the size of each node corresponds to the mean visit frequency to said decision point, and color intensity to the mean confidence index, calculated as the difference between correct and incorrect turns relative to the total number of turns per node. In Fig. 2j we visualize the values for only the optimal start to goal route (the direct path) on the binary tree. The confidence index illustrated that in the first learning trial, both sexes showed low confidence and relatively higher visit frequencies along the path, but by the memory session, the confidence increases along the route, most prominently in females, whereas males remain less confident at several intermediate junctions. To quantify these visualizations of the direct path, Fig. 2k compares memory session nodewise distributions to the random-walker performance. In the memory session, female confidence in turns on the direct path from the goal to the reward differs from the random walker (p=0.0260), whereas male confidence does not (p=0.4740). Although overall confidence or visit-count distributions of the direct path to the reward did not differ between the sexes. Overall, NaviGraph’s topological behavior metrics and visualizations yield a nuanced and process-level perspective on how spatial learning and memory unfold over time.

### Topological metrics uncover ApoE4-related memory deficiencies

NaviGraph’s topological behavior analysis was applied to a well-established humanized mouse model of the most prevalent genetic risk factor for Alzheimer’s disease (AD), Apolipoprotein E ε4 (apoE4) ^31^ which typically shows a mild behavioral phenotype ^32,33^ (Fig. 3a). Male and female apoE4 mice were compared to control mice expressing the apoE3 isoform, which is considered the benign variant associated with normal cognitive function ^31,32^. ApoE4 and control apoE3 mice underwent a learning session (four rewarded trials) followed by a memory session 72 hours later (single rewarded trial), as described in the previous section. To assess general exploration patterns, we first examined maze coverage (Fig. 3b,c), which quantifies the percentage of the maze explored over normalized session time. All mice demonstrated broader maze exploration during the learning session compared to the memory session (Wilcoxon signed rank; p < 0.02 for all groups), indicating increased task familiarity and more focused navigation during the latter. Female apoE4 mice exhibited higher maze coverage during both the learning and memory sessions, with particularly strong differences relative to apoE3 males, apoE3 females and apoE4 males during the memory session (Mann-Whitney U; p = 0.002, p = 0.014 and p=0.037, respectively).

**Figure 3:**
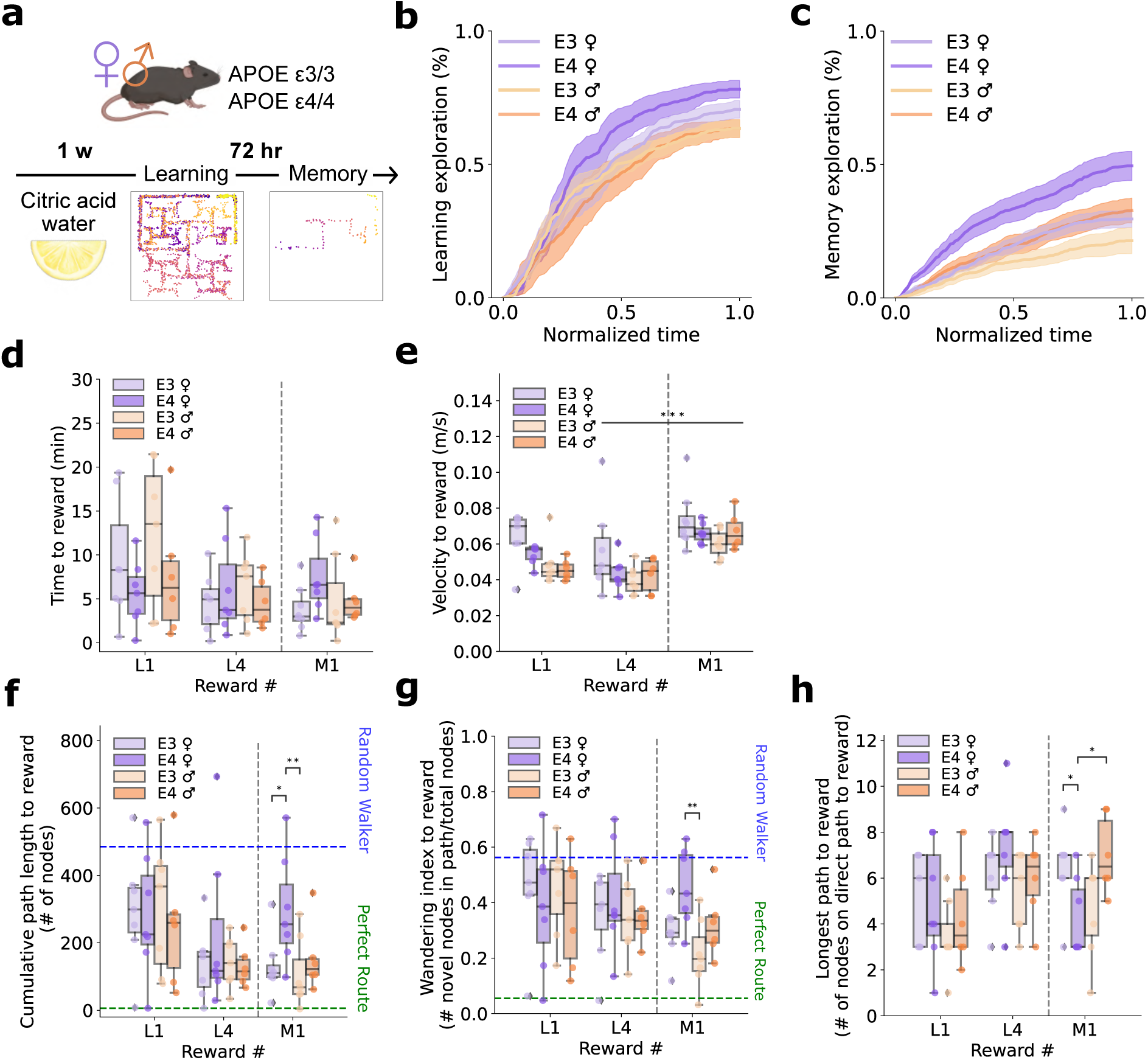
NaviGraph topological analysis uncovers behavioral nuances in an apoE4 Alzheimer’s disease model. (a) Experimental design. (b, c) Maze exploration. Mean percentage of maze explored over time, normalized to either Learning (b) or Memory (c) session durations. (d-h) NaviGraph behavioral classic and topological metrics: (d) Time to reward. (e) Average velocity to the reward. (f) Cumulative path length, total number of nodes passed to reach reward. Mean of 5000 random walker bouts on the graph depicted by the blue dashed line. Direct path to reward depicted in green dashed line. (g) Wandering index, proportion of unique nodes visited. (h) Longest direct path to reward, number of graph nodes passed on the direct path to the reward. Plots depict results for first and last rewards in learning session (L1, L4) and reward in memory session (M1). Females and males shown in purple and orange, respectively, apoE4 genotype in darker shades. Data depicted as median ± interquartile range for apoE4 and control apoE3 targeted replacement 5-month-old female (n=7-8) and male (n=6-7) mice; * p<0.05, ** p<0.01, *** p<0.001.

To reduce visual complexity in the displayed results and emphasize the differences from the beginning to the end of the learning session, only the first and final learning trials are presented in the main behavioral metrics of Figure 3, while full trial results are shown in Supplementary Fig. 1. Conventional behavioral metrics such as the time taken to reach the reward or the average velocity (Fig. 3d,e), did not reveal significant genotype or sex differences across trials. In the time to reward (Fig. 3d), a trend was observed where female apoE4 mice took longer to reach the reward compared to control apoE3 mice in the memory session, but these differences were not statistically significant. Notably, all groups displayed significantly higher velocity during the memory session compared to the final trial of the learning session (p=0.0003 for trial number; mixed-effects model analysis between the L4 and M1 rewards) (Fig. 3e).

In contrast to conventional measures, topological metrics captured within-session changes, as well as phenotypic differences. Specifically, the cumulative path length decreased significantly between L1 and L4 (p = 0.007; Mixed-effects model fixed effect for trial number) (Fig. 3f), alongside more direct trajectories (p = 0.009) (Fig. 3h). While no significant genotype or sex effects were detected within the learning phase; apoE4 females exhibited a significant increase in direct path length during the memory session compared to the final learning trial (p=0.006; Tukey’s post hoc), suggesting impaired spatial memory. In contrast, apoE3 mice and apoE4 males did not show significant differences, indicating more preserved memory in these groups. Additionally, comparisons between L1 and M1 showed residual improvement across sessions in all topological navigation measures (fixed effect for the session in a Mixed-effects model of L1 vs. M1; p=0.009 for cumulative path length, p=0.036 for wandering index, p=0.033 for direct path length).

Topological metrics further captured group-specific differences in memory performance during the recall session (M1), particularly highlighting deficits in apoE4 female mice. The longest path to the reward, which represents the sequence of correct decisions to the goal (Fig. 3h), differed significantly between groups (p=0.041, Kruskal-Wallis test). ApoE4 females followed less direct routes to the reward compared to apoE3 females (p=0.023) and apoE4 males (p=0.026), suggesting impaired recall of the maze structure. A similar pattern emerged in the cumulative path length (Fig. 3f), with apoE4 females visiting significantly more nodes on the route to the reward than apoE3 females (p=0.05) and apoE4 males (p=0.005), indicating reduced navigational efficiency. The wandering index (Fig. 3g), which reflects the extent of exploratory behavior by the proportion of unique nodes visited, further supported these findings. During the memory session, apoE4 females exhibited wandering indices approaching those observed in random walk simulations (see previous section), in contrast to apoE3 males (p=0.0025) that showed a trend approaching significance levels relative to apoE3 females (p=0.055), both of which showed more focused navigation. Overall, significant differences were observed between groups in this measure during the memory session (p=0.025), reinforcing the presence of genotype and sex-specific apoE4 memory impairments in topological navigation behavior.

To further characterize these behavioral patterns, we visualized decision confidence alongside visit frequencies (Fig. 4). As in the previous section, the confidence index was defined as the difference between mean correct and incorrect turns, normalized by total turns from each node. For each mouse, the individual confidence index at every node was computed and these values were then averaged within group and condition. We examined how confidence values scaled with visit frequency per node using linear regression. Leaf nodes were excluded from this analysis, as by definition turns from dead ends always lead back toward the reward, yielding a confidence index of 1.

**Figure 4:**
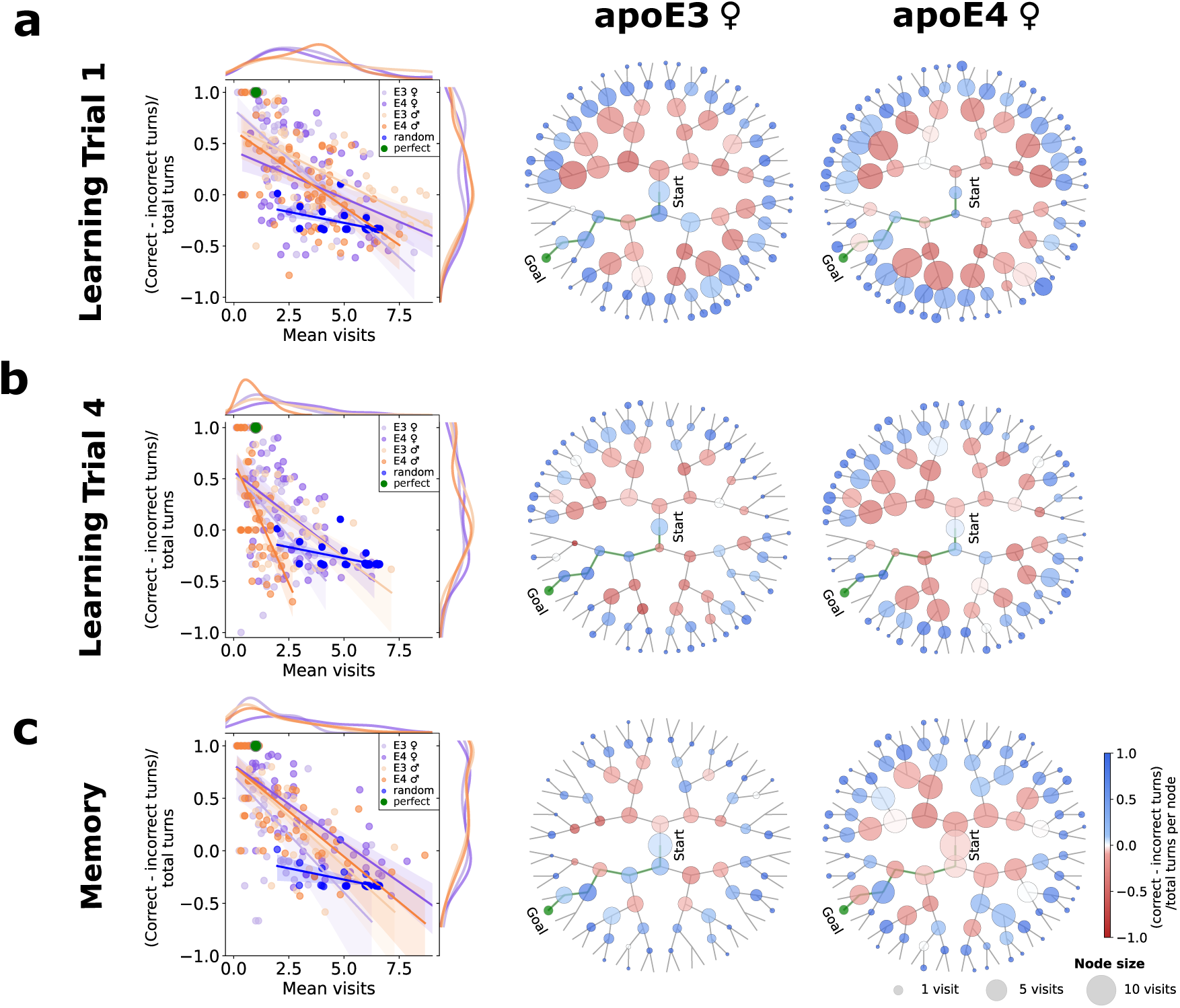
Topological analysis of navigation confidence and visit frequencies in apoE mice. (a) First learning trial. (b) Final learning trial. (c) Memory session. Left panel: Node decision confidence and visit frequency distribution relationships. Confidence index is calculated as the difference between mean correct and incorrect turns relative to the total turns made from each node. Steeper negative regression slopes indicate less confidence with increased visits. Females and males are shown in purple and orange, respectively, with apoE4 genotype in darker shades. Mean of 5000 random walker bouts depicted in blue, perfect path to reward depicted in green. Middle and right panels: Confidence index and visits per node displayed on the binary tree representing the maze, for apoE3 (middle panel) and apoE4 (right panel) female mice. Node size corresponds to the mean number of visits and color intensity reflects decision confidence. Data from apoE4 and control apoE3 targeted replacement 5-month-old female (n=7-8) and male (n=6-7) mice.

In the first learning trial (Fig. 4a), visit counts and confidence values did not differ between groups. Regression analysis revealed modest negative slopes across all groups, indicating reduced confidence at more frequently visited nodes. ApoE3 females (-0.1925) and apoE4 males (-0.1488) showed steeper declines, whereas apoE3 males (-0.1093) and apoE4 females (-0.0918) had more gradual slopes. The random walk simulation was nearly flat (-0.0428), possibly reflecting that the relationship between node visits and confidence is specific to task engagement.

By the final learning trial (Fig. 4b), visit patterns diverged significantly. ApoE4 females exhibited higher visit counts compared to apoE3 females (p=0.0004; Kolmogorov-Smirnov) and apoE4 males (p<0.0001), while differences with apoE3 males did not reach significance (p=0.080). Confidence values remained comparable across groups, yet regression slopes highlighted variation in how confidence related to visit counts. ApoE4 males displayed the steepest decline (-0.4766), apoE3 females an intermediate decline (-0.2568), and both apoE3 males (-0.1594) and apoE4 females (-0.1667) showed more moderate reductions. Random walks again yielded the shallowest slope (-0.0428), and their distributions were distinct from all biological groups. In the memory session as well (Fig. 4c), apoE4 females showed significantly higher visit counts compared to apoE4 males (p=0.006), apoE3 females (p = 0.0002), and apoE3 males (p=0.0025; Kolmogorov-Smirnov). Similar to the final learning session, regression slopes indicate a consistent reduction in confidence index with increasing visit counts across all groups. ApoE3 females (-0.2217) and apoE3 males (-0.1898) had the steepest slopes, apoE4 males were intermediate (-0.1727), and apoE4 females showed the most gradual decline (-0.1486). Interestingly, in both the final learning session and the memory session, apoE4 females displayed relatively gradual slopes. Overall, these patterns complement the behavioral differences previously observed in apoE4 female mice through topological measures, underscoring the ability of graph-based analysis to reveal subtle cognitive alterations.

### Topological neuronal, physiological and behavioral data integration

NaviGraph’s ability to integrate behavioral, neuronal and physiological data is exemplified in Figure 5, showcasing the potential to analyze these within a graph-based framework. This topological alignment enables node- and edge-specific neuronal summaries, repeated-path comparisons, and multimodal coupling within a single standardized structure. This constitutes a shift from event-triggered neural analysis toward a process-level representation embedded in the task’s decision architecture.

**Figure 5:**
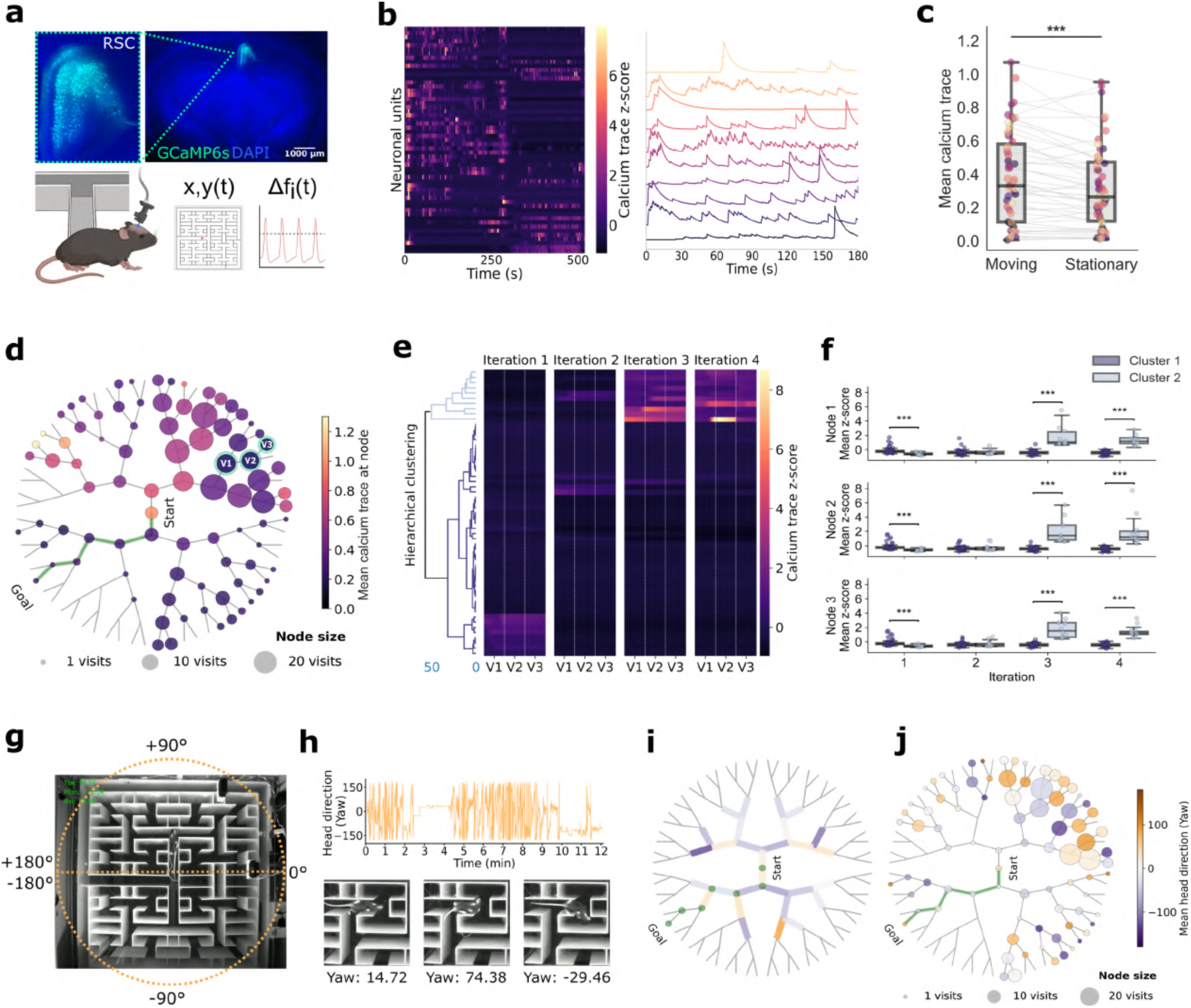
Topological integration of behavioral, neuronal and physiological data. (a) Experimental setup for simultaneous recording of calcium signals in the retrosplenial cortex (RSC) with miniature microscopes during complex maze exploration ^38^. Top panel: Coronal section of the AAV1.Syn.GCaMP6s injection site in the RSC; GCaMP6s labeled neurons (green), tissue stained with DAPI (blue). (b) Population (left) and representative individual neuron (right) activity z-score over time during maze navigation. (c) Mean calcium trace of individual neurons during movement versus stationary episodes. (d) Graph representation of the binary tree maze structure. Nodes represent decision points, node size proportional to visit frequency and color intensity reflects mean calcium trace. (e) Activity heatmaps for repeated navigations in a specific path, depicted as nodes V1, V2 and V3 in (d). Hierarchical clustering (Ward’s method) highlights distinct neuron clusters through activity patterns. (f) Neuronal cluster subpopulation activity during repeated navigation bouts to the same path (Cluster 1 n=45, Cluster 2 n=10). Boxplot data shown as median ± interquartile range; *** p<0.001. (g) Maze structure with overlaid head direction reference angles. (h) Time-series of head orientation (yaw) throughout maze exploration, with representative directions depicted in the bottom panel. (i, j) Graph representation of mean head direction at edges (i) or nodes (j), reflected by color intensity, with node size representing visit frequency.

During exploration of the complex maze, calcium signals and head direction angles were recorded with a miniaturized microscope ^25,34^ placed over the retrosplenial cortex (RSC) (Fig. 5a), an area extensively involved in spatial cognition and navigation ^35–37^. We showcase the pipeline’s ability to map this multimodal data onto behavioral decision points. This approach allows each decision point to function as a stable analytical unit; thus, neuronal fluctuations can be compared across equivalent navigational states - such as repeated arrivals to the same node - even when the animal reaches those nodes at different times, speeds, or motivational states.

A general heatmap of neuronal activity over time as well as representative neurons (Fig. 5b) demonstrate how neural activity can be tracked throughout the navigation of the maze, capturing fluctuations across different decision points. A comparison of population activity between movement and stationary periods, indicates an increase in neuronal activation during movement (p = 0.00015; paired t-test) (Fig. 5c). Further, the direct mapping of neuronal activity onto the nodes and edges of the graph structure can be combined with behavioral metrics. For example, in Figure 5d node size corresponds to visit frequency and color intensity reflects the average neuronal activity (shown as the temporal component of the calcium trace). This graph-based approach offers a straightforward way to both analyze and visualize how specific maze locations elicit varying degrees of neuronal activity. A unique advantage of the graph-based representation is the ability to isolate repeated executions of the same path, even when separated in time, and compare neuronal patterns across such repetitions. This enables a form of “topological trial averaging” that cannot be easily achieved with time-series approaches, because visits on the same route occur at irregular intervals. Using this feature, we extracted neuronal activity during multiple traversals of a defined path and applied hierarchical clustering to assess path-specific consistency (Ward’s method) on z-scored calcium temporal component activity to uncover natural groupings of neuronal activity patterns across visits to the same nodes (termed V1, V2, V3) (Fig. 5e). The resulting dendrogram provides a visual arrangement of clusters based on similarity in activity patterns over the course of the animal’s separate visits to the same path components (nodes in this case). Two dominant clusters emerged and were subsequently compared across nodes and iterations (Fig. 5f). Statistical testing (Mann–Whitney U with Bonferroni correction) showed that cluster differences varied across iterations rather than nodes. Significant differences between Cluster 1 (n = 45) and Cluster 2 (n = 10) were observed across all nodes in the first iteration (V1: p = 0.0004, V2: p = 0.0005, V3: p = 0.0005), while no differences were detected at any node during the second iteration. In contrast, iterations 3 and 4 showed divergence between clusters at all nodes, with lower activity in Cluster 1 and stronger responses in Cluster 2 (all p < 0.0001). These results point to evolving differences in activity patterns between neuronal subpopulations during repeated navigation along a defined path, possibly suggesting a functional heterogeneity within the RSC that may be further explored in the future. These illustrate the type of analysis made tractable by NaviGraph: the potential to identify functional subpopulations of neurons based on repeated topological motifs in behavior. Importantly, this is achieved without requiring rigid trial structure or explicit task epochs, highlighting the method’s relevance for naturalistic and unconstrained navigation.

Because all data streams are linked to the same graph scaffold, NaviGraph can express these as additional layers on the same nodes and edges, enabling multimodal comparisons anchored to behavioral structure rather than time. For example, we incorporate head orientation metrics. Here, yaw angles were extracted from head orientation data (see Methods section) and mapped onto the graph representation of the maze spatial coordinates. We demonstrate topological representations of mean head orientation along distinct paths (Fig. 5i) and decision points (Fig. 5j), to exemplify how mouse orientation data can align with neuronal activity and behavioral choices, providing a rather holistic view of goal-directed and exploratory behaviors. Overall, we presented NaviGraph’s ability to integrate and visualize multimodal data - including behavior, neuronal activity, and physiological parameters - into a unified analytical framework. This approach offers a novel perspective for tracking changes in neuronal encoding and behavior over time, as well as revealing patterns of plasticity and adaptation as the structure of the environment is being learned and later retrieved.

## Discussion

We introduced here NaviGraph, a flexible graph-based analysis pipeline designed to capture the intricacies of spatial decision-making and behavior across a wide range of scenario layouts. This approach responds to two ongoing challenges in behavioral neuroscience: the tendency to analyze behavior as isolated, time-point events, and the fragmentation of multimodal data across separate analytical pipelines. Conventional analysis often treats actions like zone entries or reward collection as discrete outputs, while neural or physiological signals are typically aligned to these events without considering the task’s structural context. NaviGraph addresses these by modeling spatial tasks as graphs, where decision points form nodes and transitions form edges, embedding behavioral, neuronal, and physiological data streams within a unified topological framework. This enables behavior to be interpreted as structured paths, offering a more nuanced perspective on cognition. In addition, the pipeline was designed with usability in mind. Many existing frameworks generate outputs that require extensive post-processing and custom scripting to extract group-level biological insights. NaviGraph streamlines this process, offering an end-to-end modular workflow that produces interpretable results, while minimizing the need for advanced computational expertise. Further, as NaviGraph incorporates all the data streams, behavioral setup layout and analysis results into a single data structure, sharing experimental results becomes more transparent and straightforward paving the road for standardization and improved reproducibility between laboratories.

We applied NaviGraph to both wild-type and Alzheimer’s disease (AD) risk factor apoE4 knock-in mice and demonstrated its ability to uncover subtle phenotypes not captured by conventional metrics alone. By reinforcing classic behavioral metrics such as time and velocity with topological measures, NaviGraph provides a powerful platform for studying both healthy and pathological behaviors. We also demonstrated the integration of neuronal calcium imaging and head orientation dynamics with behavioral trajectories, highlighting NaviGraph’s ability to unify diverse data within a structured spatial context.

### Graph-based behavioral analysis

To demonstrate the full capabilities of NaviGraph’s topological approach, we developed a trial-based and automated version of the naturalistic complex maze introduced by Rosenberg et al. ^8^, which is implicitly abstracted as a graph, and tracked mice behavior during a learning session and spatial memory recall session 72 hours later. This design, analyzed through NaviGraph’s synergistic combination of classic and graph-based topological metrics, allowed us to dissect behavioral changes in the complex maze across trials and between sessions. We interpreted spatial learning and task acquisition as the improved progression of behavior across the learning session trials. Spatial memory, in contrast, was assessed by comparing the final learning trial to the memory trial. If performance in the memory trial remained consistent with the final learning trial, we inferred successful memory retention. In contrast, a decline in performance was taken as indicative of impaired memory recall.

In WT mice, we validated this paradigm by demonstrating a clear learning effect across trials, indicating progressive acquisition of the maze structure and improved efficiency of decision-making over time. Although this cohort was small and primarily intended to showcase NaviGraph’s capabilities, the analysis highlighted sex-specific navigation patterns, particularly during the memory phase. While both sexes explored the maze extensively and similarly during the learning session, males displayed slightly higher maze coverage in the memory session, reflecting less efficient recall (Fig. 2). Spatial memory retention was topologically quantified using the sequence of consecutive correct decisions made to reach the reward. The longer the direct path to the reward, the fewer mistakes were made, a result interpreted as enhanced spatial memory ^8,28^. In the memory session, females followed longer uninterrupted direct paths to the reward with fewer deviations compared to males (Fig. 2i), suggesting a more accurate recall of the maze structure. Notably, these experiments were conducted in the dark, minimizing reliance on visual cues. This may have resulted in greater dependence on egocentric navigation strategies, which rely on self-movement and proximal cues ^29,39,40^. Previous reports suggest that males tend to use allocentric strategies during navigation and exploration, relying on external cues to form cognitive spatial maps of their environment, whereas females often rely on egocentric strategies ^41,42^. This could, in part, explain the more efficient navigation to the reward observed in females during the recall session, as females adapt more flexibly to tasks when external cues are limited ^43^.

To contextualize performance within topological measures, one needs to contrast performance against a random-walker agent. NaviGraph has a dedicated module to perform such simulation on any arbitrary graph that represent the task at hand. With this module, we generated benchmarks for random navigation through averaging 5000 random walks on the maze tree graph to the reward location, both with and without backtracking. Interestingly, non-backtracking random walks, which prevent revisiting nodes unless a dead end is reached, more closely resembled the navigation patterns we observed in mice. These values were subsequently used as performance benchmarks in our analysis. This observation aligns with well-established findings that animals often avoid revisiting previously explored locations to optimize navigation efficiency ^8,29,44,45^, a behavior indicative of forward planning and effective exploration driven by hippocampal mechanisms that support goal-directed planning and optimal path encoding ^46,47^. Such characteristics are reflected more accurately by non-backtracking models. This demonstrates NaviGraph’s ability to seamlessly compare real-world behaviors to computational models on the same graph, allowing for the natural identification of navigation patterns and deviations. For example, in WT male and female mice, performance during the first learning trial (L1), as measured by the total number of nodes visited and the number of unique nodes traversed per trial, appeared comparable to the random walk benchmark, consistent with the first exposure to the maze and a lack of prior spatial knowledge. By the final learning trial (L4), both measures showed a visible shift toward values more closely resembling a direct, efficient route, suggesting a transition from exploratory to goal-directed behavior over the course of the learning session.

An additional innovation enabled by the topological approach to behavior is the ability to visualize behavioral patterns directly on the maze’s graph structure. This capability provides a visually intuitive and detailed representation of navigation strategies, decision-making, and efficiency, offering insights to the experimenter that extend beyond traditional summary metrics (Fig. 2j,k). We computed a “confidence index” measure to quantify decision-making accuracy at each node within the maze relative to the reward location. This index was calculated as the difference between correct and incorrect turns from a given decision point, normalized by the total number of turns taken at that node. For instance, a node with a confidence index approaching 1 indicates highly consistent correct decision-making, while values near 0 suggest random choice behavior, and -1 portrays consistent incorrect decisions. This metric allows pinpointing specific areas where navigation falters or excels, offering a fine-grained visual analysis of behavior with both spatial and temporal precision. In this experiment, female WT mice displayed greater decision-making confidence and less node visitation relative to males in nodes leading to the reward during the memory session. The continued exploration observed in male memory sessions may reflect less effective strategy adaptation or learning under these conditions. These patterns demonstrate the approach’s sensitivity, aligning with the notion that topological structure, representing a sequential set of choices, plays a key role in spatial decision-making ^48^.

Building on this, we applied NaviGraph to a disease model using apoE4 knock-in mice, carrying the most prevalent genetic risk factor for AD ^31^. In this model, a mild behavioral phenotype and heightened sensitivity to stress is usually exhibited, limiting the efficacy of traditional assays and measures in detecting apoE4-mediated cognitive decline ^32,33,49^, specifically in young and middle-aged mice ^50,51^. We compared 5-month-old male and female apoE4 mice to control mice expressing the apoE3 isoform, which is considered to be the benign variant associated with normal cognitive function ^31^. Despite a lack of significant differences in classic metrics such as time taken to reach the reward, NaviGraph revealed differences in topological metrics of apoE4 females (Fig. 3), aligning with a large body of evidence indicating that female apoE4 carriers are at a significantly higher risk for cognitive decline and AD compared to males ^52,53^. Specifically, we observed less efficient paths to the reward location, higher wandering indices, and lower decision-making confidence during the memory session of apoE4 females. In contrast, apoE4 males demonstrated relatively preserved navigation abilities, performing more similarly to apoE3 mice across several metrics. These findings are in agreement with previous studies showing that apoE4 males are often less affected by spatial memory impairments than females, which show increased vulnerability to AD-related neuropathology, including amyloid-beta plaque accumulation and synaptic dysfunction ^50,53,54^.

NaviGraph further captures group-specific patterns by populating the graph nodes with diverse behavioral parameters such as the confidence index and visit frequencies (Fig. 4). ApoE4 females exhibited the highest visit counts during the memory session, significantly differing from apoE4 males and both apoE3 control groups. Interestingly, uniformly negative regression slopes across all groups were observed, suggesting a shared behavioral pattern: confidence is decreased in nodes where visit frequency is higher. However, the differences in the steepness of these slopes between groups highlight varying degrees of sensitivity to this dynamic, with apoE4 females usually exhibiting more gradual decline. The increased decision point visits observed in apoE4 females could partly explain their less steep regression slope, as a higher number of visits inherently dilutes the contribution of each individual turn (correct or incorrect) to the confidence index calculation, leading to a less dramatic decline in confidence as visit counts increase. Overall, our results are consistent with the finding that apoE4 females exhibit more pronounced deficits in spatial navigation. Interestingly, these findings diverge from the WT group, where females outperform males in navigation efficiency. While these findings are based on a small sample, it may reflect the broader observation that the apoE4 allele exacerbates cognitive decline in females more than in males, potentially altering sex-based differences observed in healthy mice.

Taken together, these results underscore the advantage of ecologically relevant tasks combined with a topological analysis approach to detect subtle cognitive phenotypes and highlight the potential of NaviGraph as a tool for revealing impairments in disease models where traditional behavioral measurements alone may fall short, such as apoE4 knock-in mice ^32,33^. This framework supports high-resolution behavioral analysis under minimal conditions, including limited trials and small cohorts, without imposing constraints on either. Its flexibility accommodates both short-session designs, critical for models with reduced task tolerance, and extended protocols suited for longitudinal studies, including those involving neuronal recordings such as calcium imaging or electrophysiology.

### Towards standardization of neuronal and physiological data integration

A key component of NaviGraph is its ability to incorporate additional data streams, such as neuronal activity and physiological data, to the same graph-based framework. Here, we showcased the integration of behavioral, neuronal calcium imaging and physiological head orientation data during maze exploration. Neuronal activity was imaged via miniaturized miniscopes ^25^ in the retrosplenial cortex (RSC), which has been shown to play a critical role in path integration and spatial navigation, particularly in tasks involving topological ^36^ and egocentric strategies lacking visual cues ^55,56^. We demonstrated the feasibility of mapping neuronal activity and head direction angles to specific decision points within the maze, translating these into nodes and edges on the graph for correlation with behavioral data (Fig. 5). We observed fluctuations in calcium transients across nodes, enabling a granular view of neuronal activity during spatial decision-making. Specifically, we demonstrated the ability to track this across repeated visits to the same paths, to study how neuronal activity or physiological parameters can shift with experience and familiarity. For example, hierarchical clustering revealed subpopulations which displayed varying activity relative to path familiarity, showing the potential of this approach for future comparison of neuronal activity across different behavioral states, such as exploration and goal-directed behavior. Moreover, NaviGraph provides solid ground for detecting “insight moments” or shifts in strategy ^8,28^, where neuronal activity aligns with a more efficient or learned path. Such findings could help elucidate how different cognitive states, such as task engagement or spatial learning, emerge at the neural level in health and disease.

### Future directions and conclusions

We demonstrate NaviGraph’s potential for uncovering subtle behavioral, neuronal and physiological phenotypes, however this study primarily aims to showcase our approach concept and potential use cases. While we integrate a limited sample of neuronal, behavioral and head orientation data from the retrosplenial cortex, future studies could apply the pipeline to larger datasets and additional brain regions involved in key processes like decision-making and memory consolidation ^57^. Moreover, additional data streams such as physiological or metabolic readouts could be naturally incorporated to this framework, enabling a more holistic approach. NaviGraph allows for applications beyond rodent models, with potential extensions to any experiment involving decisions in a predefined spatial layout such as virtual reality arenas used in human cognitive neuroscience. Flexible design enables future studies to leverage it across various tasks and imaging techniques, such as integrating data from mini 2-photon microscopes ^15^, or electrophysiological recordings ^58,59^.

In summary, we introduce a graph-based topological framework as a powerful and complementary approach to classical behavioral analysis of spatial decision-making. NaviGraph is a flexible, open-source tool that enables comprehensive exploration of multiple layers of information making it ideal for sharing data in a standardized fashion. By integrating diverse data streams - behavioral, neuronal, and physiological - within a unified graph structure, NaviGraph offers a high-resolution, biologically meaningful lens through which to study behavior. This framework enhances the sensitivity and interpretability of behavioral analysis and provides a novel, scalable platform for uncovering cognitive dynamics and deficits in both health and disease.

## Methods

### Mice

All procedures were approved by the Tel-Aviv University Animal Care Committee (permit number 04-20-060), and every effort was made to minimize animal stress and usage. Animals were housed under standard vivarium conditions; 22°C±1°C, 12-hour reverse light/dark cycle allowing behavioral testing during their subjective night, with *ad libitum* food and water (see Behavior measurements section). General behavioral assessments were conducted on 3-month-old C57BL/6 mice (n = 4 males, n = 5 females). For Alzheimer’s model experiments, we used apoE targeted replacement mice, in which the endogenous murine APOE gene was replaced by human apoE3 or apoE4. These mice were obtained from Taconic Laboratories (Germantown, NY) and were homozygous for the ε3 or ε4 alleles, as previously described ^60,61^. Genotypes were confirmed by polymerase chain reaction (PCR) analysis. Behavioral testing was performed on 5-month-old apoE mice (n = 7-8 females, n = 6-7 males). Simultaneous behavioral and neuronal data were collected from a single 8-month-old female.

### Behavior measurements

We developed and calibrated an automated, rewarded trial-based complex maze task based on the design of Rosenberg et al. ^8^. This maze is optimized to produce a strong spatial learning task with minimized human intervention. The paradigm requires mice to make six correct binary decisions to access a liquid reward. The reward consists of ∼5 µL of a 10% sucrose water solution administered to the reward port via an automated syringe pump ^62^ (see supporting data and Supplementary Fig. 2). Two Arduino-based automated doors with infrared (IR) movement sensors ensure unidirectional navigation post-reward to the start of the maze, allowing the next trial to begin immediately without extra handling (see supporting data for further details). Open test tubes containing 10% sucrose solution were strategically placed around all sides of the maze as a control to eliminate odor cue-based navigation. Between trials, the maze was thoroughly cleaned with 70% ethanol to ensure no residual odor cues. The maze was rotated 180° between learning and memory sessions to prevent the use of spatial cues. All behavioral testing was executed during the mice’s subjective night under IR illumination, in a self-contained light-free chamber designed for this apparatus (Supplementary Fig. 2). Behavioral imaging for the apoE experiments was acquired via Thorcam software, while general and neuronally synchronized experiments utilized a Blackfly S FLIR camera.

To motivate the mice to perform the task and minimize stress, an alternative method to water restriction was used ^26^; 1 week prior to behavioral testing home-cage water was replaced with water containing 1% food-grade citric acid (CA) (LD Carlson). This method has been previously shown to maintain stable weight and health without impacting behavioral performance ^26,27^. Mice had *ad libitum* access to CA water, and body weights were monitored daily from the onset of exposure through the completion of behavioral testing. Mice were handled and monitored by the behavior testing experimenter. Behavioral testing involved two independent sessions. In the learning session, each mouse was introduced to the apparatus through the reward arm receiving a baseline reward, followed by four rewarded trials or a maximum of 90 minutes. 72 hours later a memory session was conducted. Mice were allowed one rewarded trial or a maximum of 30 minutes. Mice that did not complete the full set of session trials within the allotted time were excluded from further analysis.

### Viral injections and cranial windows

Viral vector injections and cranial window surgeries were conducted as previously described ^63^. Briefly, mice were anesthetized with isoflurane (5% for induction, 1.5% for maintenance) and given dexamethasone (2 mg/kg intramuscularly) and carprofen (5 mg/kg intraperitoneally) for inflammation control and analgesia. Mice were positioned in a stereotaxic frame (Kopf), with body temperature maintained at 37°C. A 3 mm craniotomy was performed over the RSC of the right hemisphere, centered at 2.5 mm posterior to bregma and 0.5 mm lateral to the midline, using a high-speed manual drill (Osada, drill bit 005 Gebr. Brasseler), with intermittent cooling by artificial cerebrospinal fluid (ACSF). An adeno-associated viral vector expressing GCaMP6s (Syn.GCaMP6s.WPRE.SV40, AV-1-PV2824; titer: ∼3.6×10^12^; UPenn Vector Core) was diluted (1:2 ratio) with ACSF and injected into the RSC at two depths (-450 µm and -250 µm from the pial surface) using a glass micropipette (1 mm outer diameter and a 0.5 mm inner diameter (BF100-50-10, Sutter Instrument) pulled using a micropipette puller (P-2000, Sutter Instrument) to a final tip thickness of 20 to 45 μm). Approximately 400 nL of viral solution was delivered at each depth with pulses of 20 ms at a pressure of 20 psi using a PicoSpritzer II (Parker Hannifin). The micropipette was left in place for 10 minutes post-injection to allow for diffusion and prevent backflow, and 5 minutes between injection depths. A 3 mm coverglass was placed above the cranial window and sealed with glue (Loctite 401) surrounded by black dental cement (Ortho-Jet). The area was then covered with silicone (Ecoflex) to prevent damage to the window. Dexamethasone and carprofen were administered for 3 days following surgery as described above.

### Miniscope Ca2+ imaging

Four weeks following the viral injections, mice were anesthetized once more as described above and a miniature microscope (UCLA V4 Miniscope, Open Ephys Production Site, RRID:SCR_021480) ^34^, attached to its aluminum baseplate, was positioned over the cranial window. The field of view was optimized by focusing on blood vessels and cellular structures. Once the view was confirmed, the baseplate was cemented into place, and the miniscope was detached for future use during imaging sessions. Following base-plate placement, mice were put on a 1% CA water regimen, as described above. Mice were switched to a reverse light cycle, handled daily and habituated to the miniscope for five days in a circular arena within the same environment as the behavioral testing. During testing, animal positions were tracked and synchronized offline with miniscope data as described in the sections below.

### Tissue Processing

Mice were euthanized via an intraperitoneal injection of pentobarbital (150 µL) and transcardially perfused with phosphate-buffered saline (PBS), followed by 4% paraformaldehyde (PFA). Brains were post-fixed in PFA at 4°C for 24 hours, and then placed in 30% sucrose for 48 hours. Frozen coronal sections (50 μm) were cut on a sliding microtome, collected serially, placed in 200 μL of cryoprotectant (containing glycerin, ethylene glycol, and 0.1 m sodium-phosphate buffer, pH 7.4), and stored at −20°C. These were processed as free-floating sections and stained with DAPI (MP Biomedicals). Nuclei visualization as well as GFP expression at the viral injection site in the RSC and accurate miniscope placement were confirmed using an Olympus slide scanner VS200 microscope with a 10 x objective.

### Data acquisition and preprocessing

In experiments involving solely behavior data acquisition - the WT behavioral imaging was recorded with a FLIR Blackfly S camera and SpinView software, while the apoE experiment was imaged using Thorcam software. Behavioral data was labeled (nose, midline top, midline middle and tail base) using a locally pre-trained DeepLabCut (RRID:SCR_021391) ^17^ network. DeepLabCut outputs pose estimation data in .h5 format, containing X and Y pixel coordinates along with likelihood scores for each labeled body part, which serves as input for the NaviGraph pipeline. For combined neuronal and behavioral imaging, behavior data was recorded at 30 fps using the FLIR setup mentioned above, triggered by and synchronized with the UCLA V4 miniscope data acquisition (DAQ) system ^34^. Calcium imaging movies were processed with the MiniAn pipeline (RRID:SCR_022601) ^64^ to extract denoised temporal activity traces (C) for individual neurons.

### Head orientation analysis

Head orientation angles, stored along miniscope data in quaternion, were transformed locally to Euler angles (yaw, pitch, roll) with the SciPy Rotation library (RRID:SCR_008058) ^65^. To prevent discontinuities at the ±180° boundary, angles were transformed into Cartesian coordinates, and then converted back for angular analysis. The yaw angle only was used for current head direction calculations. To obtain the phase offset between the sensor reference frame and the experimental reference frame, the linear equation between the positions of the 3 midline body parts of the mouse that did not include the nose was calculated for each frame (midline top, midline middle and tail-base). Logically, when all body parts tagged in this dataset are aligned the mouse should be looking straight ahead. Accordingly, we extracted the mean yaw angle for frames in which the nose was positioned along the computed linear equation when the mouse is facing the root node of the maze, resulting in the reference value. Processed yaw values were synchronized and incorporated to NaviGraph through a dedicated plugin (see pipeline section below), enabling analysis of head orientation in relation to behavior and neuronal activity at decision points and paths.

### Navigraph pipeline

NaviGraph is a modular Python-based pipeline for processing and analyzing behavioral, neuronal, and physiological data in maze paradigms using a graph-based framework (Fig. 1). Its plugin-based architecture provides complete flexibility, enabling the integration of diverse data streams, graph architectures, analysis metrics, and visualization modules. The codebase includes a command-line interface for setup, calibration, testing, and execution, as well as straightforward installation through package managers. Multiple example configurations are provided in the repository, illustrating applications across different mazes and data streams. Full installation and setup instructions can be found in the GitHub repository https://github.com/PBLab/NaviGraph.

#### Configuration

Pipeline execution is defined in YAML configuration files, which specify experimental parameters, data paths, graph properties, analysis modules, and visualization options, allowing customization for various experimental designs and data streams.

#### Calibration

Behavioral data are aligned with the maze schematics through a transformation matrix derived from user-selected reference points. This step ensures accurate spatial registration of trajectories within the experimental setup. Calibration is performed as follows:

1. Reference point selection: The user manually assigns corresponding reference points on the behavioral video and the maze map.
2. Transformation matrix computation: A transformation matrix is calculated to enable consistent spatial alignment across sessions.
3. Registration testing: To validate the calibration, newly selected points on the behavioral video are transformed using the computed matrix and overlaid onto the maze map for visual inspection.

#### Data processing and integration

Following registration, environments are segmented directly into graph-defined regions, where nodes represent decision points or areas of interest and edges denote possible transitions. Additional data streams, including neuronal activity and head orientation, are synchronized and coupled to the same graph scaffold through dedicated plugins. The pipeline generates structured datasets that integrate positional, neuronal, physiological, and user-defined inputs within a common representation.

#### Data analysis

NaviGraph facilitates the computation of both classic behavioral measurements and graph-based metrics. These are easily extended through user-defined functions. Results are aggregated and stored in a new dataset, storing all experimental sessions and corresponding behavioral metrics. The key analytical components include:

### Topological metrics

#### - Cumulative path length

Number of nodes traversed between any two points, giving a topological understanding of the animal’s movement along the maze. As a benchmark of random navigation for the purpose of this study, we used the average cumulative number of nodes on the path to reward passed by 5000 random walk bouts on the graph with backtracking allowed only when no forward walking options exist (averaged 485.38 nodes in this case) relative to the perfect direct route to the reward (6 nodes).

#### - Wandering Index

Ratio of unique nodes traversed to the total number of nodes in the graph, providing a measure of the exploratory nature of the animal’s trajectory. The random walker average (0.567) relative to the perfect route was calculated in a similar manner to the previous metric.

#### - Direct path to reward

Calculates the longest uninterrupted sequence of nodes along the direct path the animal took towards the reward. This includes allowances for a set number of permitted deviations as specified by the user, making this metric highly flexible to accommodate many forms and limitations of mouse behavior around branches of the physical maze.

#### - Average time at node

Aggregation of time spent at each decision point, divided by visit frequency, to determine average dwell time.

### Classical behavioral metrics

#### - Time and velocity analysis

Computation of transition time and average movement speed between positions in the maze.

#### - Exploration percentage

Calculation of the proportion of unique nodes visited from the total number of nodes in the graph, providing a measure of maze coverage.

### Random walk simulations

NaviGraph includes a built-in module for generating random-walk trajectories directly on any graph representation of behavioral paradigms. This module samples transitions from the adjacency structure of the graph, drawing each step with uniform probability unless weights are explicitly provided. The user may specify the start node, an optional target node, the maximum number of steps, and the number of independent walks to generate, with an optional seed ensuring reproducibility. For each walk, the pipeline records the full node sequence, enabling subsequent computation of path-length statistics and success rates when a target node is defined. In this study, non-backtracking random walks were used to construct a null model of navigation on the same graph geometry, providing a benchmark against which animal trajectories could be compared.

### Customizable analysis

The modular structure of NaviGraph allows for easy integration of additional user-defined behavioral, neuronal or physiological analysis metrics tailored to specific research questions.

### Result aggregation and visualization

Visualization modules are defined in the configuration file, including trajectory overlays which allow for synchronized visualization of behavioral, neuronal, and physiological measures across diverse experimental paradigms. Data aggregation by experimental groups and sessions enable advanced plotting in various formats such as graph-based visualizations, quantitative plots, or custom user-defined outputs.

### Statistical analysis

For two-group comparisons in NaviGraph behavioral metrics (male vs female WT mice Fig. 2c), Repeated Measures ANOVA test with Tukey’s correction for multiple comparisons was used in the learning session trials, and the Mann-Whitney U test was applied in the memory session comparisons. For multi-group comparisons (apoE3 and apoE4 across genders Fig. 3d-h), in the learning session (between L1 and L4 trials) a mixed-effects model with Tukey’s correction for multiple comparisons was used, and in the memory session the Kruskal-Wallis test followed by the two-stage step-up method of Benjamini, Krieger and Yekutieli for multiple comparison correction ^66^ was used. Maze exploration across sessions (Fig. 3b,c) was quantified using area-under-the-curve (AUC); within-group session differences were tested with paired Wilcoxon signed-rank tests, and between-group comparisons used Mann-Whitney U tests with Bonferroni correction. Linear regression and the Kolmogorov-Smirnov test were applied for confidence index and visitation frequency relationship assessment and distribution analysis, respectively (Fig. 4a, d). For comparing neuronal activity data while moving or stationary, first normality was confirmed using the Shapiro-Wilk test following which a paired t-test was used for comparisons. The Ward method was applied for hierarchical clustering of neurons with similar activity patterns in different iterations to the same path, followed by the Mann-Whitney U test with a Bonferroni correction for multiple comparisons of clusters. Statistical analysis was performed using GraphPad Prism 10 (RRID:SCR_002798), as well as Python’s SciPy (RRID:SCR_008058) ^65^ and Statsmodels (RRID:SCR_016074) ^67^ libraries, with a significance threshold set at α = 0.05.

## Author contributions

AKI, DMM and PB designed the study. AKI and PB designed and built the setup. AKI performed experiments, collected and analyzed data. EI and AKI developed the software pipeline. LB imaged additional behavioral paradigms. AKI and PB wrote the manuscript. DMM and PB provided financial support and mentoring.

## Acknowledgements

This work is dedicated to the memory of Daniel M. Michaelson, whose mentorship and partnership were integral to the development of this project. The authors would like to thank Eran Rosen, Nahum Aminov and the staff at the Mechanical Workshop of the School of Chemistry and Physics at Tel-Aviv University for manufacturing the maze, and Dr. Anton Sheinin from the Sagol School of Neuroscience for designing the maze automation hardware. We would also like to thank Dr. David Kain for valuable input regarding surgery, Rotem Yam-Shahor for imaging brain slices, and Lior Lin and Omri Koren for assisting in behavioral experiments. PB wants to thank the La Caixa Foundation (grant No. HR23-00560) and Israeli Science Foundation (grant No. 2342/21) for financial support. AKI wants to thank The Prajs-Drimmer Institute for the Development of Anti-Degenerative Drugs for scholarship support.

## Supplementary Figures

**Supplementary figure 1:**
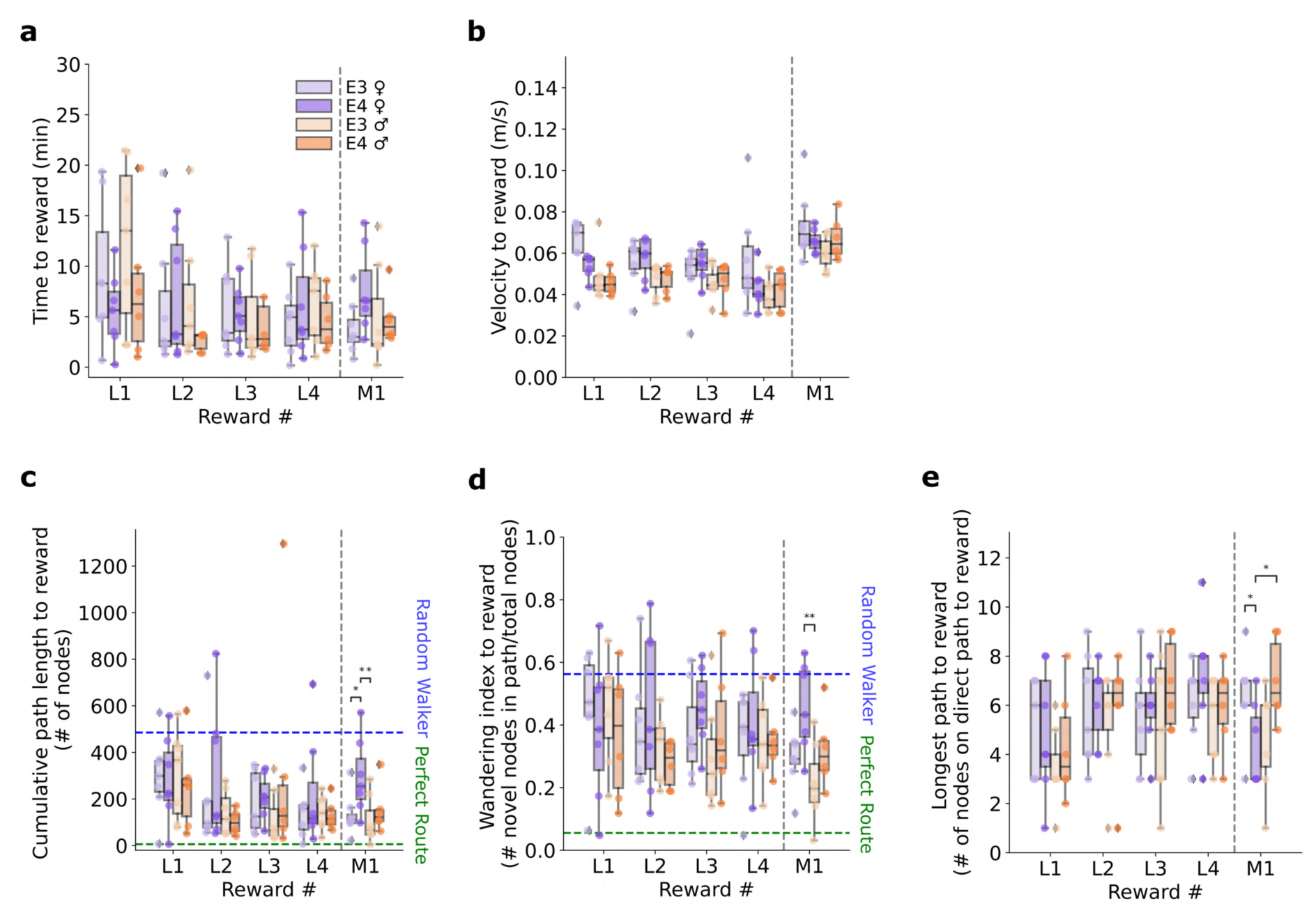
NaviGraph behavioral analysis of apoE4 and apoE3 male and female mice. ApoE4 and control apoE3 targeted replacement 5-month-old female (n=7-8) and male (n=6-7) mice received ad libitum citric acid water for one week followed by a learning and post 72h memory session in the complex maze paradigm. (a-e) NaviGraph behavioral classic and topological metrics. Plots depict results for the learning session trials (L1, L2, L3, L4) and the memory session (M1). Females and males shown in purple and orange, respectively, apoE4 genotype in darker shades. (a) Time to reward. (b) Average velocity to the reward. (c) Cumulative path length, total number of nodes passed to reach reward. Average of 5000 random walker bouts on the graph depicted by the blue dashed line. Direct path to reward depicted in green dashed line. (d) Wandering index, proportion of unique nodes passed to reach reward. (e) Longest direct path to reward, number of graph nodes passed on the direct path to the reward. Data depicted as median ± interquartile range; * p<0.05, ** p<0.01.

**Supplementary figure 2:**
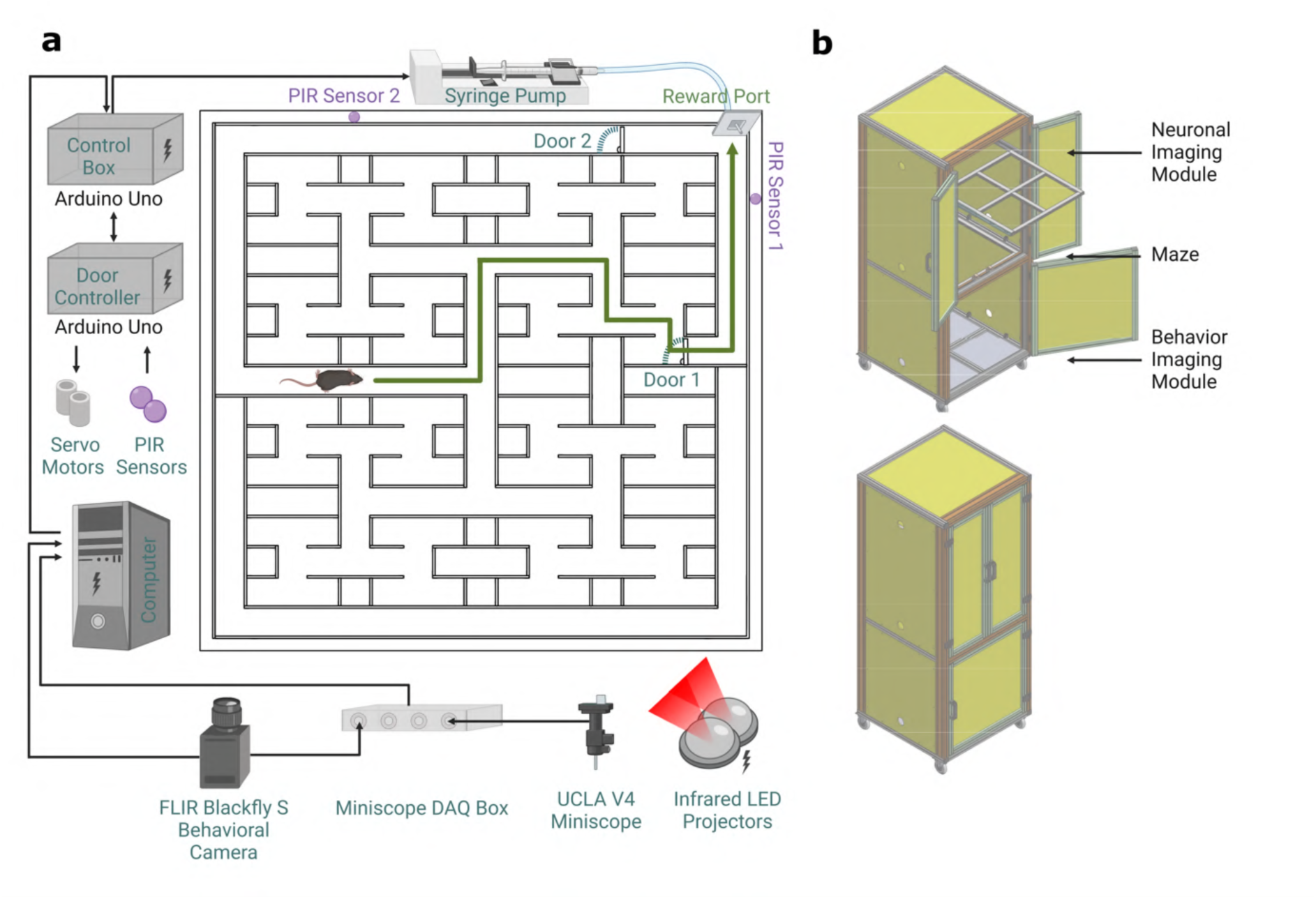
Complex maze paradigm hardware setup. (a) Schematic of the hardware components supporting the complex maze paradigm. (b) Self-contained chamber designed for behavioral and neuronal imaging of complex maze apparatus. Created with BioRender.

## Notes

https://github.com/PBLab/NaviGraph

### Competing Interest Statement

The authors have declared no competing interest.

### Summary of Updates

Updated title, erroneously shorten in first version submission. All other content is unchanged

